# Efficient High-Quality Metagenome Assembly from Long Accurate Reads using Minimizer-space de Bruijn Graphs

**DOI:** 10.1101/2023.07.07.548136

**Authors:** Gaëtan Benoit, Sébastien Raguideau, Robert James, Adam M. Phillippy, Rayan Chikhi, Christopher Quince

**Affiliations:** Organisms and Ecosystems, Earlham Institute, Norwich, NR4 7UZ, UK; Gut Microbes and Health, Quadram Institute, Norwich, NR4 7UQ, UK; Genome Informatics Section, National Human Genome Research Institute, Bethesda, MD, USA; Sequence Bioinformatics, Department of Computational Biology, Institut Pasteur, Paris, France; Warwick Medical School, University of Warwick, Coventry, CV4 7AL, UK

## Abstract

We introduce a novel metagenomics assembler for high-accuracy long reads. Our approach, implemented as metaMDBG, combines highly efficient de Bruijn graph assembly in minimizer space, with both a multi-*k*′ approach for dealing with variations in genome coverage depth and an abundance-based filtering strategy for simplifying strain complexity. The resulting algorithm is more efficient than the state-of-the-art but with better assembly results. metaMDBG was 1.5 to 12 times faster than competing assemblers and requires between one-tenth and one-thirtieth of the memory across a range of data sets. We obtained up to twice as many high-quality circularised prokaryotic metagenome assembled genomes (MAGs) on the most complex communities, and a better recovery of viruses and plasmids. metaMDBG performs particularly well for abundant organisms whilst being robust to the presence of strain diversity. The result is that for the first time it is possible to efficiently reconstruct the majority of complex communities by abundance as near-complete MAGs.

## 1 Introduction

Shotgun metagenomics, i.e. the untargeted sequencing of DNA fragments from a mixed sample of genomes in a community, is now an established tool in microbial community analysis [1]. It enables the sequencing of genetic material from microbes that cannot otherwise be studied, for example through isolation and culturing [1]. Pioneering studies have used metagenomics to survey the taxonomic and functional composition of various microbiomes such as the human gut or soil [2]. A recent large-scale reanalysis of all shotgun metatranscriptomes has enabled the discovery of an order of magnitude more RNA viruses than previously known [3]. A critical first-step in metagenomics analyses is the assembly of shotgun reads into longer contiguous sequences or contigs.

While short-read sequencing produces petabytes of valuable metagenome data each year, genome assemblies derived from short reads are typically highly fragmented into millions of contigs per sample, preventing the precise assessment of genomic contents. The low quality of assemblies is due to intra- and inter-genome sequence repeats, low coverage of some species, and strain diversity. Many complete genomes are nevertheless recovered by clustering (i.e. *binning*) short contigs using features such as sequence composition or differential coverage across multiple samples [4], creating so-called metagenome assembled genomes (MAGs). While MAGs have resulted in thousands of bacterial genomes being added to reference databases, typical MAGs from short-read metagenomes remain fragmented, contaminated, and missing key regions such as the 16S rRNA gene operon.

Third-generation long-read sequencing technologies have greatly improved the quality of metagenome assemblies and MAGs. The first applications using reads generated by the Oxford Nanopore technology, which at that point had a relatively high error rate, typically only resolved a small fraction of the community as complete circularised contigs [5]. More recent results using the long (*≈* 10 kbp) and accurate reads generated by HiFi PacBio have improved on this dramatically, with hundreds of genomes obtained as circularised contigs or partially fragmented MAGs [6].

Existing *in silico* techniques for producing MAGs using long-read metagenome assembly remain limited. Firstly, whilst assembly of complete lineage resolved MAGs is now possible, both low abundance and highly abundant organisms with strain diversity may not be assembled [7]. The result is that the majority of the community by abundance is still not resolved as high-quality MAGs [6]. Secondly, the computational performance of current tools severely limits the size of samples that can be be processed: oceanic/terrestrial samples or large co-assemblies remains prohibitive to assemble. Even typical metagenomes require long processing times (days) and high-end computing infrastructure (*>* 500 GB – 1 TB memory) which is out of reach for many labs. Thirdly, most current assemblers do not allow the easy incorporation of contextual data such as depth of coverage that is a critical component in metagenome reconstruction.

There are two generally accepted paradigms for sequence assembly, string graph methods that operate with individual reads, considering pair-wise overlaps and constructing graphs to represent these [8], and de-Bruijn graph (dBG) assemblers where reads are first decomposed into short fixed-length sequences (*k*′-mers) [9]. The former, requires all-vs-all read comparisons which scales poorly with read number, and hence, is too inefficient for short-read metagenomics. It has been applied to long reads, specifically HiFi PacBio metagenomics, in hifiasm-meta [10]. This is made possible through the use of minimizers to efficiently find overlaps between reads prior to assembly. Minimizers are a means of selecting a subsample of *k*′-mers in a reproducible way so that similar regions across reads share the same minimizers. The decomposition to *k*′-mers in dBG assemblers enables them to reduce the volume of data to process and efficiently detect overlaps. They are now the typical approach for large-scale short read data sets. To the best of our knowledge, they have not been applied to long-read metagenome assembly, apart from an hybrid approach (Flye [11]) which uses a form of sparse de Bruijn graph, termed as A-Bruijn graph [12]. With carefully subsetted *k*′-mers, it forms initially noisy disjointig assembled sequences, that are then used to create a repeat graph, further resolved through read mapping. This approach works for both noisy Nanopore and accurate HiFi PacBio sequences, and has been adapted to metagenomics, but does not scale particularly well, nor does it provide state-of-the-art performance on HiFi data [13, 6].

String graph, de-Bruijn graph, and hybrid approaches all have limitations when applied to long-read metagenome assembly. String graph methods are still relatively inefficient and coverage estimation is difficult on the uncorrected string graph because of ambiguous read mapping when strain diversity or noise generates complex graphs with many similar alternate paths. There are two challenges in applying de-Bruijn graph to long reads, firstly they effectively assume exact overlaps, and secondly for long reads the required overlap and therefore *k*′-mer size becomes large and the number of unique *k*′-mers required and hence memory, prohibitive. Scalability, will become a key issue for HiFi PacBio metagenomic assembly. This data type has the potential to allow genome resolution from even complex metagenomes such as soil and plankton, which will be transformative, but only if we can develop tools that can scale to large amounts of sequence data whilst addressing the unique characteristics of metagenomes of uneven coverage depth and strain heterogeneity.

A fundamentally different approach to the problem of adapting dBGs to long reads was introduced with rust-mdbg [14], which uses a minimizer space de-Bruijn graph (MDBG). The units of the MDBG are no longer *k*′-mers but sequences of minimizers of size *k*′ (*k*′-min-mers), each of which is a short *k*′-mer. The result is highly scalable genome assembly, just 12 million *k*′-min-mers are required to assemble a complete human genome. This approach also has the appeal that it can deal better with noise than long nucleotide *k*′-mer dBGs because exact matches are only required on the small selected minimizers. The rust-mdbg algorithm, is however, not designed for metagenomics, in particular, it cannot cope well with variable genome coverage depths.

We introduce metaMDBG, a method that takes the principle of minimizer space assembly and engineers it specifically for metagenomics, at the same time introducing a number of novel algorithmic advances. We designed a highly efficient multi-*k*′ approach, where the length of *k*′-min-mers is iteratively increased whilst feeding back the results of the last round of assembly. This enables us to deal with the variable coverage depths found in metagenomes. We implemented several techniques to estimate and refine *k*′-min-mer abundance as their length is increased. This information on abundance is then used in a novel ‘local progressive abundance filtering’ strategy to reduce the graph complexity generated by errors, inter-genomic repeats and strain variability. We demonstrate on multiple mock and real community metagenome HiFi PacBio data sets, that metaMDBG, has an order of magnitude better scaling of memory than the current state-of-the-art whilst outperforming all existing algorithms in terms of near-complete MAGs and run time.

## 2 Results

### Overview of metaMDBG

We present metaMDBG, a method for assembling metagenomes from accurate long reads (e.g. PacBio HiFi). metaMDBG takes as input a set of reads and outputs a FASTA file with contigs. The overall assembly strategy is summarised in Figure 1-A. The universal minimizers, which are *k*′-mers that map to an integer below a fixed threshold (see Methods), in each read are first identified. Each read is thus represented as an ordered list of the selected minimizers, denoted a minimizer-space read (mRead). Each iteration of the assembler then comprises the construction of a de-Bruijn graph using lists of minimizers of fixed length *k*′, denoted *k*′-min-mers, starting with 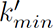 = 4. We count *k*′-min-mers across the whole dataset and those with frequency below a set threshold are filtered (Figure 1-B). The graph is then constructed and graph simplification performed. This process includes classical methods for contig generation such as tip clipping and bubble popping. This is followed by a ‘local progressive abundance filter’ method to remove potential inter-genomic repeats, strain variability, and complex error patterns (Figure 1-C). This starts by identifying long seed unitigs, i.e. long non-branching paths in the graph. We then increment an abundance threshold starting at one up until 50% of the coverage depth of this seed. At each step unitigs with coverage equal to or lower than the threshold are removed and the graph re-compacted. This strategy, coupled with techniques for refining unitig coverage estimation (Figure 1-B), enables the seed unitig to conservatively converge on its longest possible form as complexity from the graph is removed. This completes one iteration in our multi-*k*′ approach in this minimizer space. The resulting minimizer-space contigs (mContigs) are added to the set of input mReads in the next iteration and these steps repeated but after incrementing *k*′ by one. At the end of the multi-*k*′ process when 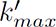, reads are mapped to the final mContigs in order to extract their base-space sequence.

**Figure 1:**
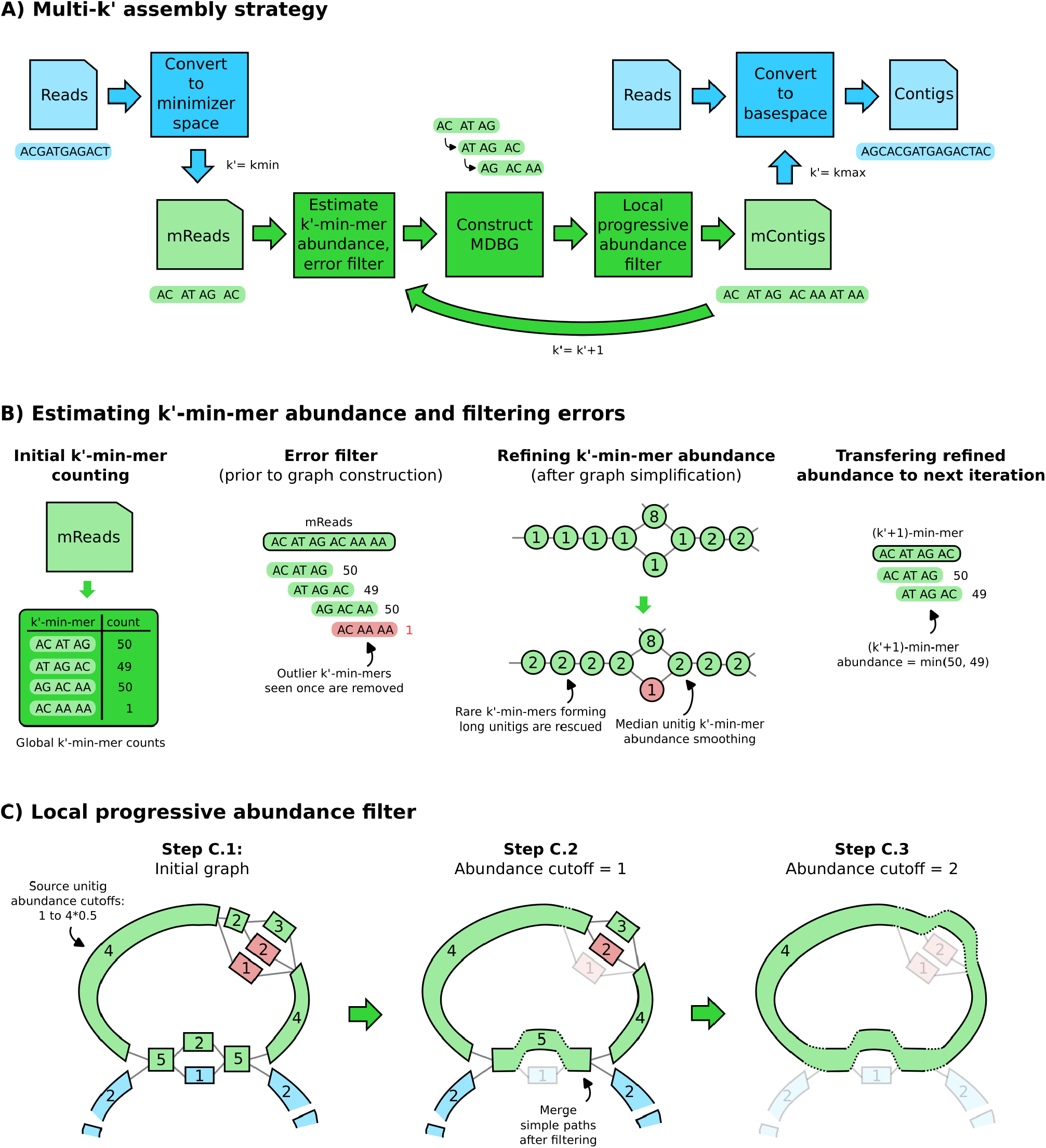
Overview of the algorithmic steps of metaMDBG. (**A**) Overview of the multi-*k*′ assembly strategy. Processes in blue are performed at the level of nucleotide sequences, while the ones in green are performed at the level of minimizers only. (**B**) Components for estimating and refining *k*′-min-mer abundance as *k*′ is increased, and filtering errors prior to graph construction. (**C**) Illustration of the ‘local progressive abundance filter’ algorithm that simplifies complex graph regions generated by errors, inter-genomic repeats and strain variability. Each node represents an unitig (unitigs in green and blue belong to two distinct species, unitigs in red represents errors). The long unitig on the top-left part of the graph is chosen as seed (step C.1). Its abundance (4) is used as reference to apply a ‘local progressive abundance filter’ from one-times to half its abundance (step C.2 and C.3). At each step, unitigs with abundance equal to the cutoff value are removed, then the graph is re-compacted to simplify fragmented unitigs. Note that fragmented green unitigs with abundance 2 would have been removed without the intermediate step C.2.

In the following sections, we demonstrate that metaMDBG accurately reconstructs bacterial genomes and outperforms state-of-the-art approaches. Furthermore, metaMDBG only uses a fraction of the computational resources needed by other tools, enabling for the first time an accurate and cost-efficient reconstruction of large metagenomes.

### Benchmarking setup

We compared metaMDBG with two other state-of-the-art assemblers for HiFi metagenomics data: metaFlye (v2.9-b1768) and hifiasm-meta (v0.2-r058). We do not include a comparison to the rust-mdbg implementation of minimiser space de Bruijn graphs [14], because it was not competitive on the metagenome data sets considered here. This is unsurprising since it is designed specifically for genome assembly.

We experimentally evaluated the assemblers on two mock communities and three real metagenomic projects from different environments (Table 1). The two mock communities, ATCC [15] and Zymo, contain respectively 20 and 21 species for which abundances and reference genomes are known. The real datasets derive from three metagenome sequencing projects of various sequencing depths and from microbiomes with different levels of species diversity. The first one ‘Human’ is a PacBio generated dataset composed of four human fecal samples from omnivore and vegan donors. The second one ‘AD’ is a time-series of three samples extracted from anaerobic digester sludge, generated for this study. For these two projects, where multiple samples were available, we present results below from coassemblies of all samples together, although results on single-sample assembly are available in Supplementary Table S2. The third dataset ‘Sheep’, is a single deeply sequenced sample from the sheep rumen [6].

**Table 1:** Evaluated metagenome datasets.

| Sample | Accessions | Number of bases (Gb) | N50 read length (kb) | Sample description |
| --- | --- | --- | --- | --- |
| ATCC | SRR11606871 | 59 | 12 | Mock community ATCC MSA-1003 [15] |
| Zymo | SRR13128014 | 18 | 10.6 | Mock community ZymoBIOMICS D6331 |
| Human | SRR15275213, SRR15275212, SRR15275211, SRR15275210 | 68 | 10.5 | Four human gut datasets from a pool of omnivore and vegan samples |
| AD | ERR10905741, ERR10905742, ERR10905743 | 178 | 10.1 | Time-series of three samples extracted from anaerobic digester sludge (week t+1, t+20 and t+40) |
| Sheep | SRR14289618 | 206 | 11.8 | Sheep rumen microbiome [6] |

Since the true genomes in the real datasets are not known, we used CheckM (v1.1.3) to evaluate the genome completeness and contamination level of each contig and determine whether they are MAGs. We grouped MAGs into three conventional categories based on the CheckM results: ‘near-complete’ if its completeness is *≥* 90% and its contamination is *≤* 5%, ‘high-quality’ if completeness *≥* 70% and contamination *≤* 10%, ‘medium quality’ if completeness *≥* 50% and contamination *≤* 10%.

### Improved recovery of complete circularized microbial genomes from metagenomes

We first evaluated the assemblers on two mock communities: ATCC and Zymo, by aligning contigs to references and computing average nucleotide identity (ANI) - (see Methods and Supplementary Table S3). All three assemblers performed similarly on the mocks, both in terms of number of species obtained as circularised contigs and ANI to reference sequences (*>* 99.99% in most cases). The mock ATCC contains 20 species, but only 15 with sufficient coverage depth for assembly, of these each assembler obtained 12 as circularised contigs, but not the same twelve, each assembler assembled one species uniquely. The Zymo mock contains 21 genomes, but five have very low coverage, and five are strains of *E. coli*. In this case, metaMDBG and hifiasm-meta both obtained ten circularised genomes, and metaFlye nine, but metaMDBG additionally generated two almost complete (*>* 99.8%) genomes as linear contigs. No assembler could correctly resolve all the *E. coli* strain diversity, metaMDBG circularised the most abundant strain B1109 and the strain B766 as a single linear contig, hifiasm-meta assembled strain B766 only as a circular contig but had all other strains present as fragmented contigs, and metaFlye produced fragmented genomes for all five *E. coli* strains.

Assembling sequences from real complex microbiomes is a substantially more challenging problem than for mock communities, due to the higher species richness, uneven coverage distributions and strain diversity. For all three real communities, we observe a significant improvement in the number of circularised near-complete MAGs longer than 1 Mb (cMAGs) generated by metaMDBG compared to the state-of-the-art (Figure 2-A). metaMDBG assembled 75 cMAGs on the Human gut microbiome dataset (13 more than hifiasm-meta), 114 on the AD dataset (61 more than hifiasm-meta) and 266 on the Sheep rumen dataset (3 more than hifiasm-meta). MetaFlye produced significantly fewer cMAGs than the other two assemblers.

**Figure 2:**
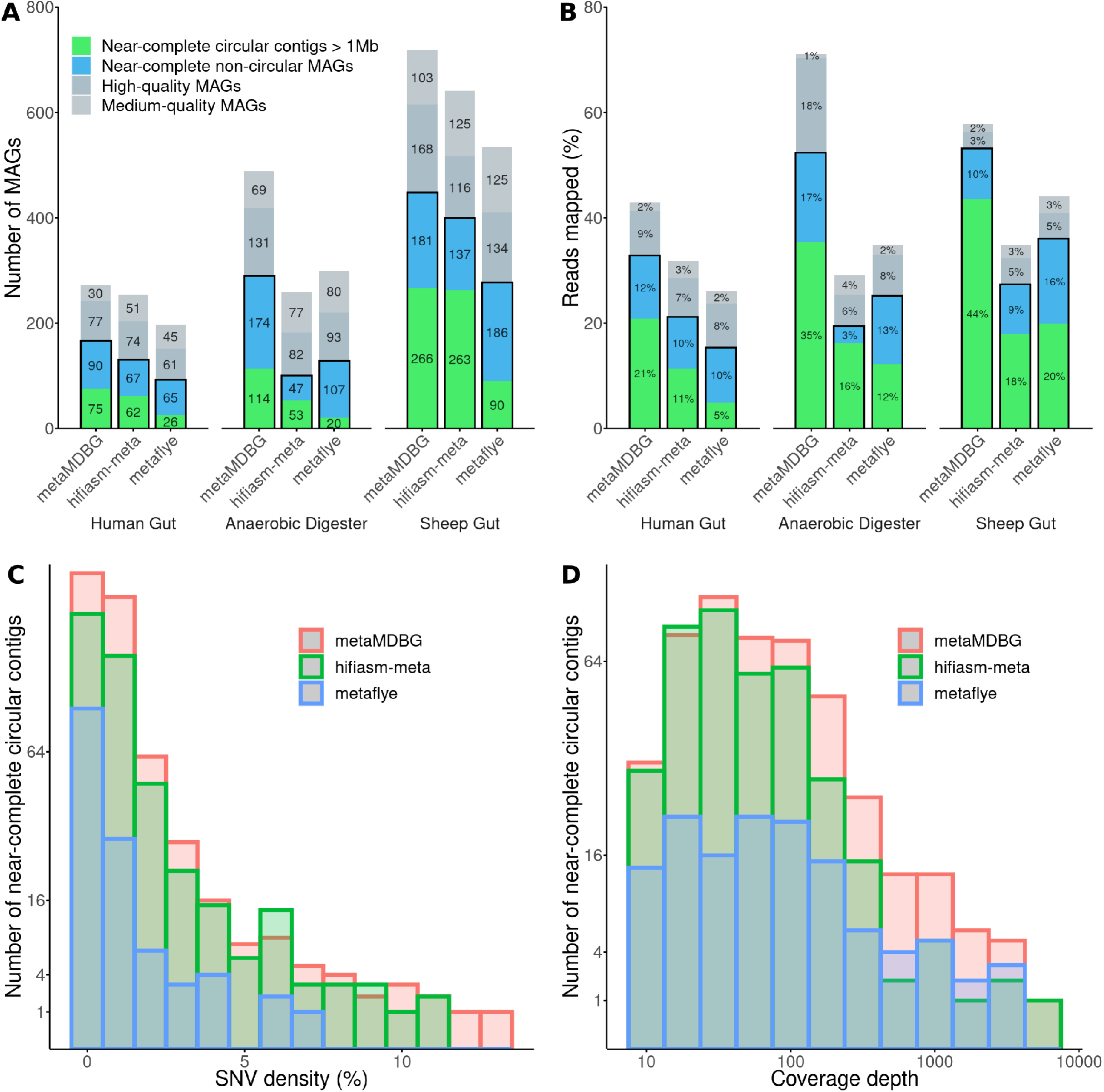
Assembly results on three metagenomic projects. ‘Human Gut’ represents the co-assembly of the four human gut samples, ‘Anaerobic Digester’ is the co-assembly of the three AD2 time-series samples. (**A**) CheckM evaluation. A MAG is ‘near-complete’ if its completeness is *≥* 90% and its contamination is *≤* 5%, ‘high-quality’ if completeness *≥* 70% and contamination *≤* 10%, ‘medium quality’ if completeness *≥* 50% and contamination *≤* 10%. (**B**) The percentage of mapped HiFi reads on MAGs. (**C-D**) The distribution of SNV density (%) and coverage depths for near-complete circular contigs generated by each assembler on all datasets (the y-axes have been sqrt-scaled); here the bars are overlaid and not stacked.

To investigate potential differences between the circular MAGs generated by the assemblers, we aligned the assemblies against each other with wfmash (Supplementary Tables S4 and S5). On the Sheep rumen dataset, metaMDBG and hifiasm-meta combined found a total of 356 distinct near-complete circular contigs. Among them, 176 where found by both assemblers (49%), with 90 specific to metaMDBG and 87 to hifiasm-meta. We note that the vast majority (91%) of these specific cMAGs are still present in the other assemblies but as one or more linear contigs. The cMAGs missed by metaMDBG were less fragmented, with a median of one contig (mean = 1.3) necessary to cover a cMAG reconstructed by another assembler, compared to a median of three (mean = 10.7) for hifiasm-meta and also a median of three for metaFlye (mean = 5 — see Figure S1). The remaining 9% of cMAGs that exist in one assembler and were not found by the other assemblers usually correspond to phased strains by hifiasm-meta or to species with low coverage that might be incomplete in other assemblies. On the human gut microbiome samples, we observed similar results in terms of specific circular contigs and linear contig fragmentation. However, on the AD dataset, the cMAGs missed by hifiasm-meta and metaFlye were highly fragmented, being covered by in both cases a median of six contigs, whereas metaMDBG still obtained a median of two contigs.

We next determined the impact of coverage depth and strain diversity, as measured by SNV density - computed with Longshot [16], on the ability of the different assemblers to resolve cMAGs from all three datasets combined. metaMDBG and hifiasm-meta were able to generate cMAGs across a range of SNV densities (Figure 2-C), but we found a highly significant negative relationship between SNV density and the probability that metaFlye assembles a cMAG (Logistion regression coeff. = -1.35, p-value = 2.18e-11, sample size *n*=575) and in fact no cMAGs with greater than 7% SNP density were found by metaFlye. For coverage depths between 10x and 100x metaMDBG and hifiasm-meta had similar success at resolving cMAGs but at higher coverages, more than twice as many cMAGs were obtained by metaMDBG (see Figure 2-D).

### Improved reconstruction of circularised phage and plasmid genomes

In addition to prokaryotic genomes, plasmids and phages will be present in a typical metagenome. These are usually smaller but can be present with high coverage depth and strain diversity presenting particular assembly challenges. We used viralVerify [17] to identify the circular components that are potentially plasmid or phage genomes (Supplementary Table S2). In all three data sets, and for both plasmid and phages, metaMDBG identified substantially more circularised plasmids and phages than either of the other two assemblers. In the Sheep rumen metagenome, metaMDBG obtained 70% more circularised plasmids and 25% more phages than the next best assembler, hifiasm-meta. In the Human gut coassembly, 42% more plasmids, and 55% more phages and in the AD coassembly more than twice as many plasmids and 55% more phages compared to hifiasm-meta which was second best in both cases. This improved performance on mobile elements by metaMDBG probably derives from the same robustness in the presence of high coverage depth and strain diversity, that we observed in the case of cMAGs.

### The majority of complex communities by abundance recovered as circular or non-circular nearcomplete MAGs

To date no existing HiFi PacBio assembler has succeeded in recovering the majority of a microbial community by abundance as near perfect MAGs from complex communities. To reconstruct noncircular MAGs, we took the contig collections for each assembler after first subtracting all circularised contigs *≥* 1 Mb, and then binned the remaining contigs with MetaBAT2, using sequence composition and coverage (across multiple samples for the Human and AD coassemblies). The contigs removed prior to binning will include the cMAGs identified above, this procedure ensures that bins are only constructed from potential genome fragments [10]. We then evaluated these bins with CheckM as described above.

The complete list of MAGs recovered by each assembler and their statistics is provided in Supplementary File S1. Figure 2-A shows that metaMDBG reconstructs 23 (34%) more near-complete non-circular MAGs than hifiasm-meta on the human gut co-assembly, 127 (270%) more on the anaerobic digester time-series and 44 (32%) more on the sheep gut dataset. MetaFlye produced less near-complete circular contigs than other assemblers, but has an equivalent or higher number of near-complete or high-quality MAGs than hifiasm-meta across all datasets, and lower or equivalent to metaMDBG. The high quality bins from all assemblers typically contained less than ten contigs (Figure S2) and MAGs that were more fragmented than this generally had low coverage.

The improvement in the number of near-complete non-circular MAGs of metaMDBG is mostly due to a better recovery of low-abundance organisms (Figure S3). This is in contrast to the cMAGs, discussed above, where metaMDBG exhibits better performance on high coverage depth genomes. The assemblers also differ in the nucleotide divergence of the near-complete MAGs they resolve. We illustrate this in Figure S4 where we show for each assembler on each data set, how the number of dereplicated near-complete MAG clusters, both circular and non-circular, collapses as they are dereplicated at decreasing levels of nucleotide similarity. In the Sheep rumen and Human gut data sets, the number of dereplicated MAG clusters from hifiasm-meta drops significantly below a 97% ANI dereplication threshold, this is not observed for metaMDBG or metaFlye, which indicates that a greater proportion of the hifiasm-meta MAG diversity is at the strain-level. This is not the case for the AD data set where no assembler seems to generate a substantial number of strains with more than 97% ANI.

To summarise the microbial diversity obtained as near-complete MAGs from the AD coassembly, we constructed a phylogenetic tree at the genus level, using a panel of single-copy core genes, for all MAGs from all assemblers (Figure 3A). This reveals that the improved MAG recovery by metaMDBG, translates into a far more representative picture of microbial diversity at all levels of evolutionary divergence. In total, we observe 114 genera that are recovered from the AD datasets by metaMDBG but missing from the near-complete MAG collections of the other programs. When the other assemblers did recover MAGs from the same genus, in all but one case metaMDBG found more MAGs, indicating a better recovery of diversity at the species level. Finally, we can see large parts of the tree in Figure 3 that are only represented by metaMDBG MAGs, in fact, 6 phyla (46 families) are only found by metaMDBG, against 1 phylum (4 families) specific to metaFlye and 4 families to hifiasm-meta. These statistics are summarised in Figure 3B and a detailed list of taxa recovered by only one of the assemblers given in Supplementary File S3.

**Figure 3:**
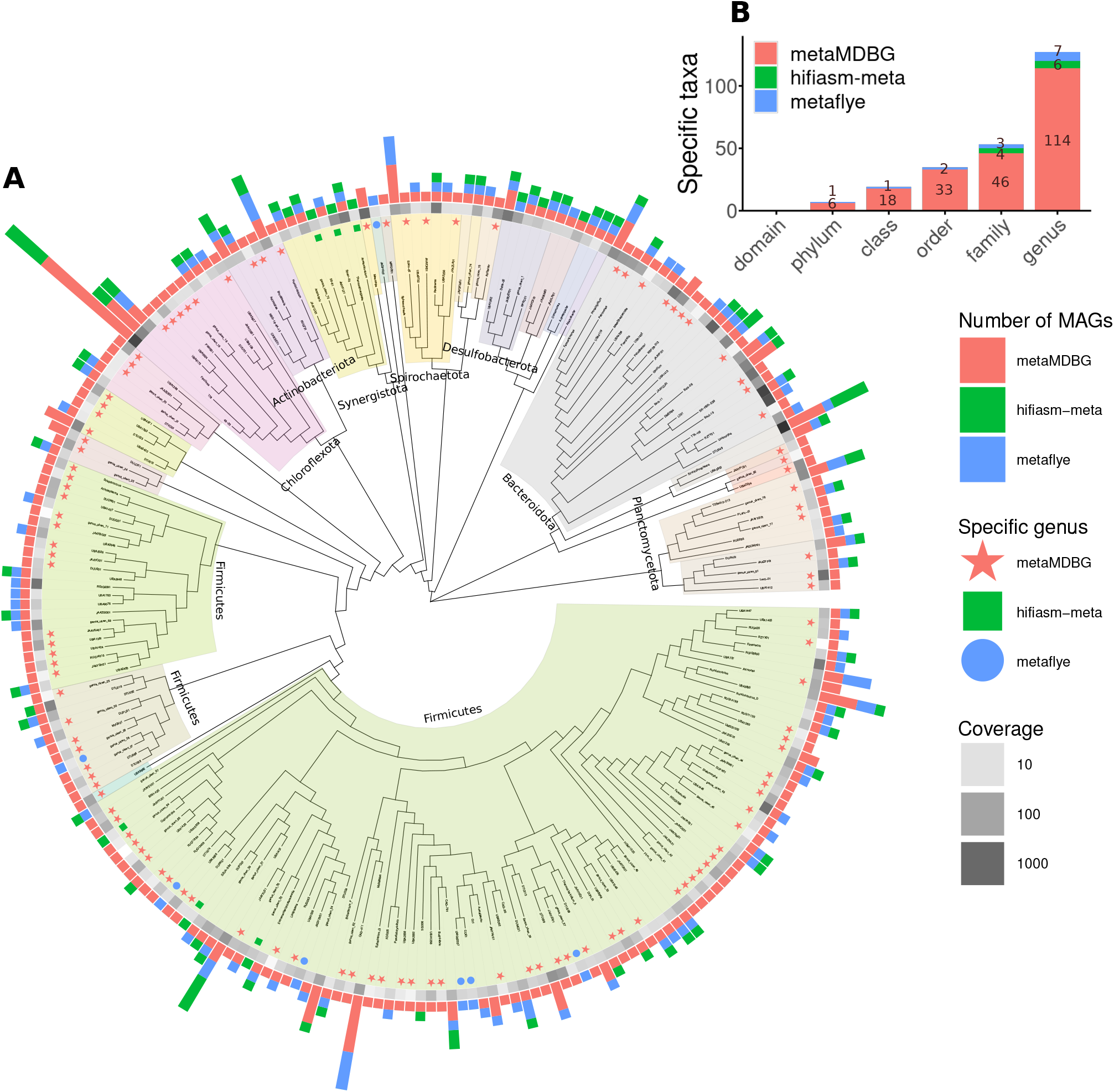
Phylogenetic tree of genera recovered from the AD dataset for all assemblers combined. (**A**) For the near-complete bacterial MAGs, we generated a *de novo* phylogenetic tree based on GTDB-Tk marker genes and display at the genus level. The outer bar-charts give the number of MAGs found in each genus. The coloured symbols then denote genera recovered by only one of the assemblers. The grayscale heat-map denotes the aggregate abundance of dereplicated MAGs in a genus. (**B**) Number of taxa at different levels that are unique to each assembler.

The combined result of these differences: the higher number of abundant cMAGs; the better resolution of low coverage near-complete non-circular MAGs; and near-complete MAGs that are more representative of the phylogenetic diversity present; is that for the AD and Sheep data sets metaMDBG actually succeeds in obtaining a collection of near-complete MAGs that can map over 50% of the reads in the original samples (Figure 2-B). This is highly significant as it implies that the majority of the community by the abundance is present as highquality constructs. This was not the case for the Human data set which may be to due to the relatively lower depth of sequencing of these samples. Considering, cMAGs alone, across all data sets metaMDBG, recruited twice as many reads as either metaFlye or hifiasm-meta (Figure 2-B).

### Efficient large-scale assembly with a substantial reduction in memory footprint

MetaMDBG is highly scalable, both in terms of execution time and memory footprint (Table 2). On the Human dataset, metaMDBG took 36h to complete, which is 20% faster than other assemblers. This gain increased significantly on more complex Sheep and AD datasets. MetaMDBG took about 3 days to assemble the AD datasets against 8 days for metaFlye and 39 days for hifiasm-meta. We observed similar trend on the Sheep dataset. With regards to memory usage, metaMDBG required only 14 GB to assemble the Human dataset, while metaFlye and hifiasm-meta used more than 130 GB. The memory consumption of metaMDBG on the AD and Sheep samples only spiked at 16 GB and 22 GB despite the larger diversity detected in those datasets. On the other hand, metaFlye and hifiasm-meta memory usage was many times this, MetaFlye required about 450 GB to complete and 650 to 800 GB for hifiasm-meta.

**Table 2:** Performance on the three metagenomic projects. Assemblers wall-clock time using 16 cores and memory footprint. These were run on a 192 core Linux x86-64 server running Intel(R) Xeon(R) CPUs E7-8850 v2 @ 2.30GHz.

|  | Human Gut (68 Gb) |  |  | Anaerobic Digester (178 Gb) |  |  | Sheep Gut (206 Gb) |  |  |
| --- | --- | --- | --- | --- | --- | --- | --- | --- | --- |
|  | meta-MDBG | hifiasm-meta | meta-flye | meta-MDBG | hifiasm-meta | meta-flye | meta-MDBG | hifiasm-meta | meta-flye |
| Time | **36 h** | 42 h | 43 h | **77 h** | 950 h | 187 h | **108 h** | 587 h | 153 h |
| Memory | **14 GB** | 163 GB | 137 GB | **16 GB** | 669 GB | 458 GB | **22 GB** | 811 GB | 439 GB |

## 3 Discussion

We have introduced metaMDBG, an assembler for long and accurate metagenomics reads based on the minimizer-space de-Bruijn graph. The principal motivation for using this structure was to develop a scalable assembler, in this we succeeded, metaMDBG on a range of data sets, was 1.5 to 12 times faster than the state-of-the-art and required between one-tenth and one-thirtieth of the memory. Moreover, we achieved this with substantially better assembly results particularly in strain-diverse communities such as the AD dataset and for the first time succeeded in reconstructing the majority of communities by abundance as near-complete MAGs. We also demonstrated improved results for phages and plasmids. This was made possible thanks to a specialized multi-*k*′ strategy for assembling rare and abundant species, coupled with a novel method for removing sequence complexity based on organism abundance.

The improvement in running time is significant when we consider the time-scale necessary to process metagenomics projects which can be counted in days. The improvement in terms of memory footprint was even more substantial, we could assemble the largest data sets with one-twentieth of the requirements of our competitors. This is important, for smaller scale projects it is democratising, enabling research groups without access to sophisticated HPC to easily assemble their data sets, even on laptops. At the other end of the scale, the potential of long accurate reads and probable future technological advances that will reduce the cost to generate them, means that we are likely to see ever larger high accuracy long read projects studying highly complex environments such as soil, metaMDBG is uniquely placed to address those assembly challenges.

Even though metaMDBG was able to reconstruct more MAGs than other assemblers, we have seen that they do not necessarily collect the same organisms. For instance, on the sheep rumen dataset, metaMDBG and hifiasm-meta found respectively 266 and 263 near-complete circular contigs, but after dereplication, only 176 were common to both assemblers. This highlights the complementarity of both methods and the possibility for future methodological improvements, e.g. through assembly reconciliation [18].

We have demonstrated that none of the assemblers directly phase strains reliably. Our approach, metaMDBG, reconstructs more contigs with high SNV density than its competitors (Figure 2), the other assemblers struggle to reconstruct consensus-level assemblies in high-diversity regions, but we do not attempt, here, to resolve finer scale variation beyond this consensus. However, we believe that these contigs could form a framework for strain phasing within the same core data structures that we have introduced. Ekim *et al.* [19] have recently developed a read mapping strategy in minimizer space. This could be used for fast mapping of reads to the assembly graphs and combined with a method for distinguishing noise from true strain variation, would enable phasing of SNPs within strains. This would be similar in principle to the approach of Strainberry [20] but integrated directly into the assembly process. That the MDBG structure preserves information on strain variability and accessory genomes is demonstrated by the observation that Ekim *et al.* [14] were able to build and query a massive MDBG pangenome from 661,405 bacterial species.

A similar approach, but based on mapping Hi-C contact reads onto the minimizer-space graphs, might also enable strain-resolution or linkage of plasmids to genomes [6]. The key point is the flexibility of our data structures to incorporate and manipulate additional information on the graph. Which we believe we enable further extensions of the underlying algorithms.

In summary, we have demonstrated the power of minimizer-space de Bruijn graphs, for assembly of highly accurate long reads from metagenomes. Outperforming all existing assemblers in terms of results and computational efficiency. We believe that further advances of our methodology coupled to larger data sets will go a long way to finally achieving complete genome-scale resolution of even complex metagenomes.

## Acknowledgements

C.Q., G.B. and S.R. are supported by the Core strategic Program of the Earlham Institute BB/CCG1720/1. S.R. is also funded through NERC grant ResPharm (NE/T013230/1). C.Q. was also partially funded by MRC Methodology Grant ‘Strain resolved metagenomics for medical microbiology’ MR/S037195/1. This project has received funding from the European Union’s Horizon 2020 research and innovation programme under the Marie Sk-lodowska-Curie grants agreements No. 872539 and 956229 (R.C.). R.C. was supported by ANR Transipedia, SeqDigger, Inception and PRAIRIE grants (ANR-18-CE45-0020, ANR-19-CE45-0008, PIA/ANR16-CONV-0005, ANR-19-P3IA-0001). AMP was supported by the Intramural Research Program of the National Human Genome Research Institute, National Institutes of Health. We acknowledge the NIH Intramural Sequencing Center (NISC) for assistance with PacBio sequencing. We would also like to thank Dr Sergey Nurk for stimulating discussions during the course of this study and Dr Orkun Soyer for supervising collection of the AD samples through the BBSRC project BB/N023285/1.

## Author Contributions Statemen

G.B. devised and implemented the approach. G.B. and S.R. performed the experimental evaluation. R.J. prepared DNA extracts for sequencing. A.M.P. and C.Q. coordinated sequencing of the AD samples. G.B. R.C., C.Q. conceived the study, supervised and coordinated the work. All authors wrote, reviewed, edited and approved the manuscript.

## 6 Supplementary Figures

**Figure S1:**
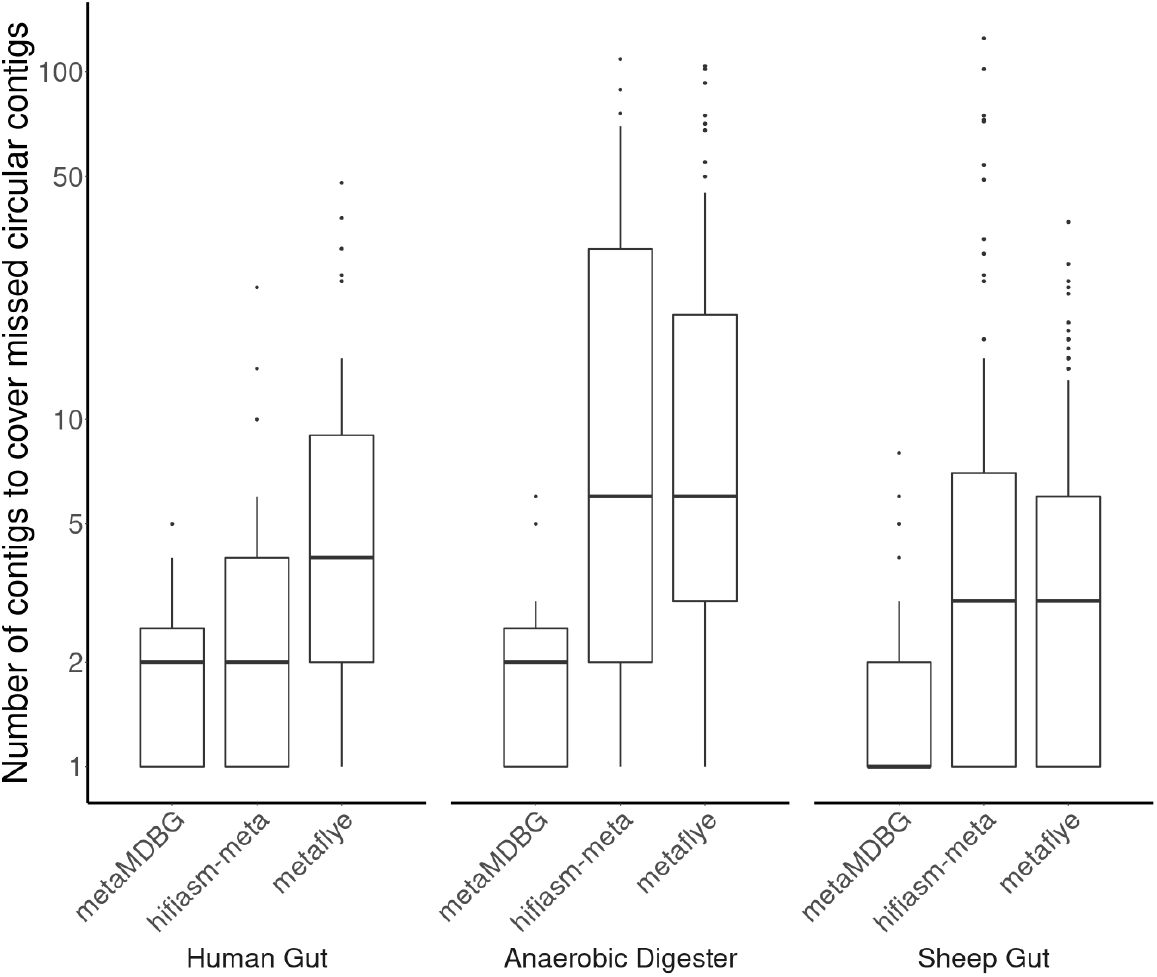
Number of contigs required to cover a near-complete circular MAG reconstructed successfully by an alternative assembler. In order to estimate the degree of fragmentation of assemblers, we aligned the contigs of one assembler against the near-complete circular contigs (cMAGs) recovered by the other assemblers. The fragmentation is then represented as the number of contigs required to cover these cMAGs (see section ‘Assessment of completeness and fragmentation of assemblies using reference sequences’ for details). The boxplot elements are the median (horizontal bar), 25th and 75th percentiles (box limits Q1 and Q3), Q1-1.5*IQR and Q3+1.5*IQR (whiskers, IQR=Q3-Q1) and outliers. Summary statistics (min, median, mean, max): Human— metaMDBG (1, 2, 2.1, 5); hifiasm-meta (1, 2, 4, 24); metaFlye (1, 4, 7.5, 48) : AD—metaMDBG (1, 2, 2.3, 6); hifiasm-meta (1, 6, 19.8, 109); metaFlye (1, 6, 15.1, 104) : Sheep— metaMDBG (1, 1, 1.8, 8); hifiasm-meta (1, 3, 10.7, 125); metaFlye (1, 3, 5, 37). The data to generate this boxplot have been extracted from the columns ‘AssemblyStatus’ of Supplementary Table S4.

**Figure S2:**
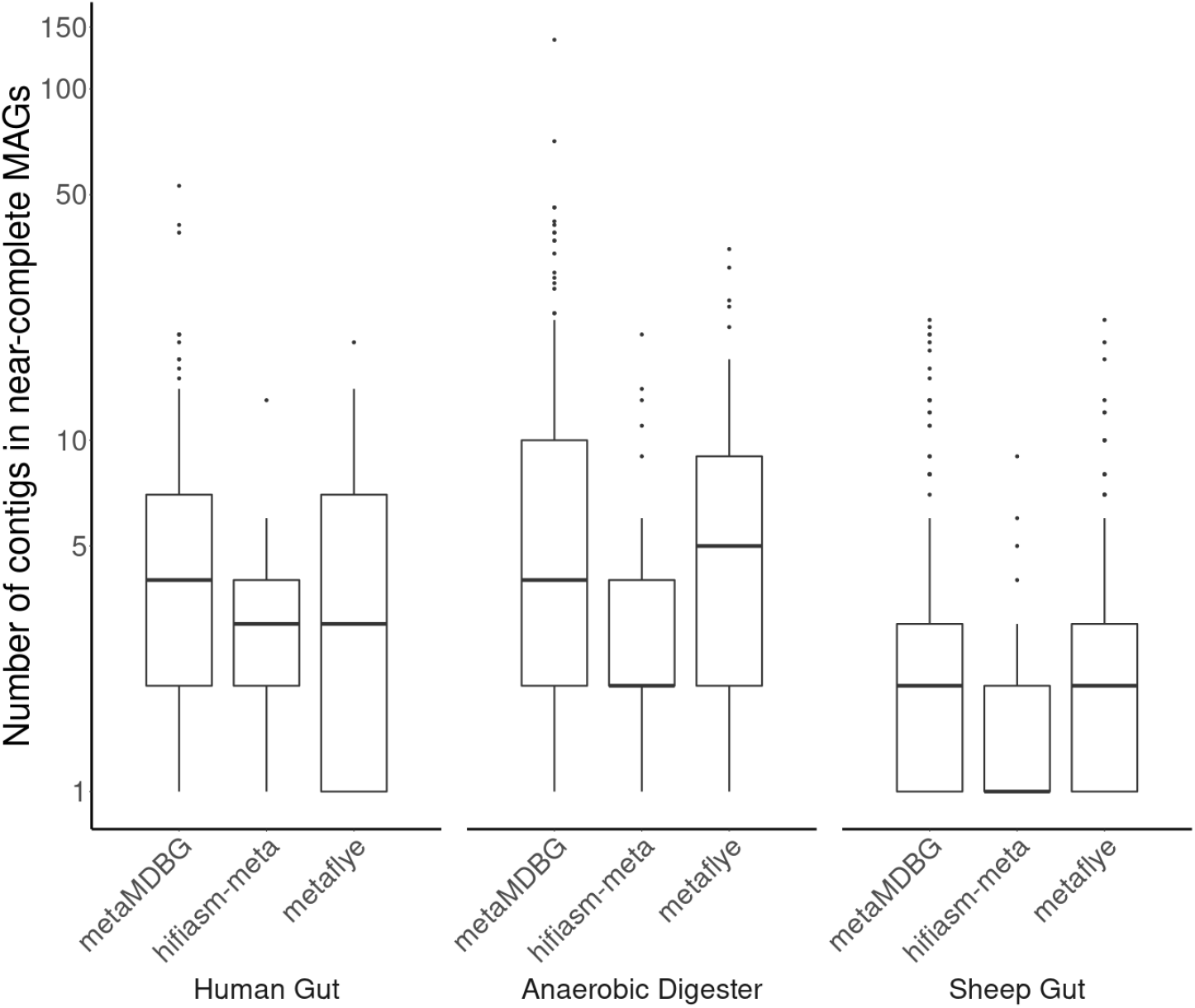
Number of contigs in non-circular near-complete MAGs. The boxplot elements are the median (horizontal bar), 25th and 75th percentiles (box limits Q1 and Q3), Q1-1.5*IQR and Q3+1.5*IQR (whiskers, IQR=Q3-Q1) and outliers. Summary statistics (min, median, mean, max): Human— metaMDBG (1, 4, 6.8, 53); hifiasm-meta (1, 3, 3.1, 13); metaFlye (1, 3, 4.6, 19) : AD— metaMDBG (1, 4, 9.2, 138); hifiasm-meta (1, 2, 3.6, 20); metaFlye (1, 5, 7, 35) : Sheep— metaMDBG (1, 2, 3.3, 22); hifiasm-meta (1, 1, 1.5, 9); metaFlye (1, 2, 2.8, 22). The number of contigs per non-circular MAGs have been extracted from columns ‘AssemblyStatus’ of supplementary File S1.

**Figure S3:**
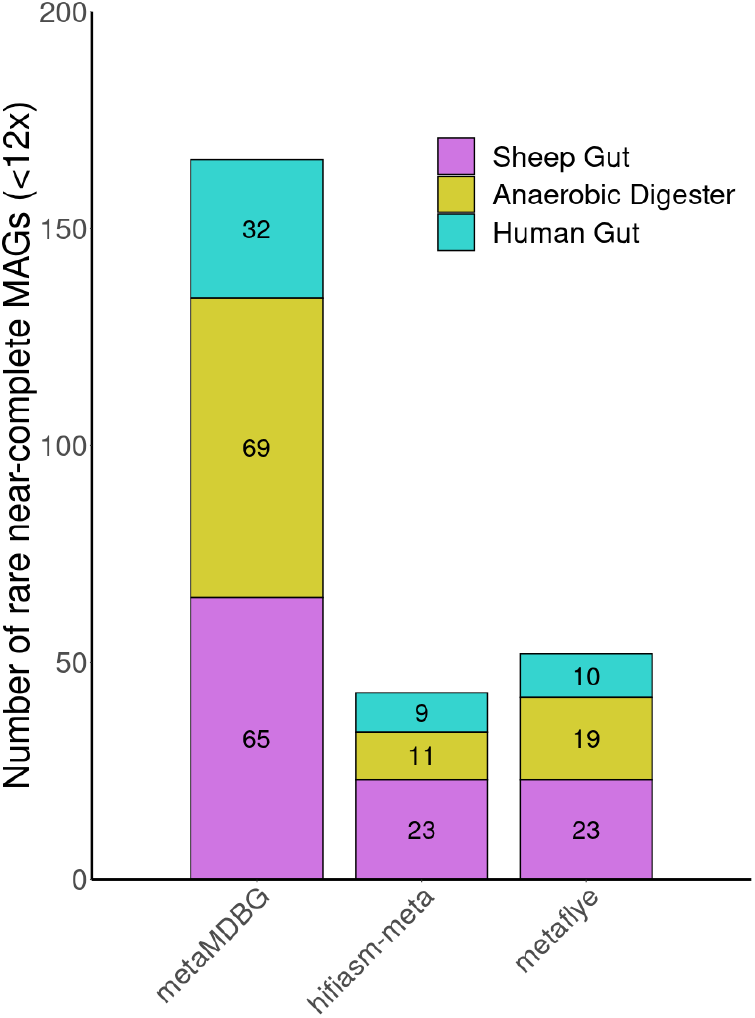
Number of low-coverage non-circular near-complete MAGs recovered by the assemblers. For the three tested datasets, we show the number of non-circular near-complete MAGs with low coverage (*<* 12*x*) reconstructed by each assembler.

**Figure S4:**
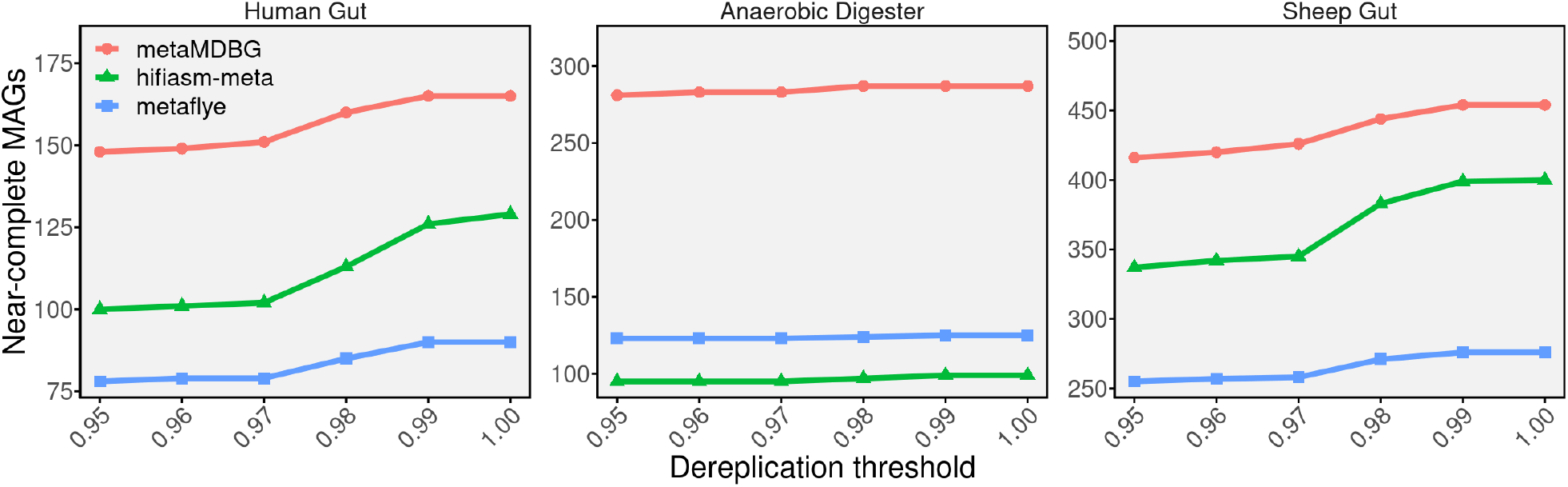
Total number of near-complete MAGs (circular and non-circular) across different dereplication thresholds. We used dRep [21] to cluster MAGs by nucleotide similarity using the parameter -sa from 0.95 to 1.

## 7 Methods

### 7.1 Preliminaries

We start with a lexicon of some terms and concepts related to minimizer-space de-Bruijn graphs (MDBG) and genome assembly.

#### Minimize

In this work, we consider the concept of universal minimizer as in [14]. Recall that in the original definition of minimizers [22], a window is used to compute minimizers. Universal minimizers are pre-determined and do not require a window to be defined. Specifically let *f* be a function that takes as input a *k*′-mer (string of size *k*) and outputs an integer value within range [0*, H*[where *H* is typically equal to 2^64^. Given 0 *< d <* 1 and k *>* 0, a universal minimizer is any string *m* of length *k* over the DNA alphabet such that *f* (*m*) *< d.H*. The value of *d* represents the density of *k*′-mers that will be considered as minimizers over the space of all possible *k*′-mers.

#### Minimizer-space read

Prior to minimizer-space de-Bruijn graph construction, each read is scanned and its minimizers are identified. Each read is therefore represented as an ordered list of minimizers. We call this minimizer representation of a read the minimizer-space read, or *mRead*.

#### *k*′-min-mer

A *k*′-min-mer is a list of *k*′ successive minimizers. They are collected by sliding a window of size *k*′ over the mReads.

#### Minimizer-space de Bruijn graph

The minimizer-space de-Bruijn graph (MDBG) is constructed from the set of *k*′-min-mers. A MDBG is a directed graph where nodes are *k*′-min-mers and an edge exists between two nodes *x* and *y* if the suffix of *x* of size k’−1 (*i.e.* its k’−1 first minimizers) is equal to the prefix of *y* of size k’−1 (*i.e.* its k’−1 last minimizers). We defer details about reverse-complementation to the ‘Assembler implementation details’ section.

#### Unitig

An unitig (or simple path), is a maximal-length sequence of distinct nodes in the graph such that, given a unitig length *n*, (1) for each 1 *≤ i ≤ n −* 1, in and out-degrees are equal to 1, (2) if *n >* 1, the out-degree of *u*_0_ is 1 and the in-degree of *u_n_* is 1. Singleton nodes (*n* = 1) are also considered to be unitigs.

#### Unitig abundance

We define the unitig abundance as the median abundance of its constituent *k*′-min-mers.

#### Minimizer-space contig

Contigs have the same definition as unitigs, they are unitigs obtained after graph simplification. Contigs are first extracted as ordered lists of *k*′-min-mers (a path in the graph). The minimizer-space representation of a contig, called a *mContig*, is constructed by concatenating the first k’ − 1 minimizers of its first *k*′-min-mer and the last minimizer of each following *k*′-min-mer (*i.e* the sequence of *k*′-min-mers without their k’−1 overlapping region). The mContig representation will be used to extract (k’+1)-min-mers in the multi-*k*′ algorithm.

#### Contig

At the end of the assembly process, the mContigs are converted to base-space by concatenating the base-space sequence spanned by the minimizers (see section ‘Converting to base-space and assembly postprocessing’ for more details).

### 7.2 Algorithmic components

The overall assembly workflow is given in Figure 1. Input reads are first converted into their minimizer-space representation (mReads). We then initiate a multi-*k*′ assembly algorithm, in minimizer space. The following operations are performed during each iteration. The abundance of *k*′-min-mers is determined and low-abundance *k*′-min-mers, deemed as erroneous, are discarded. A MDBG graph is then constructed and classical assembly graph simplification steps, such as tip clipping and bubble popping, are performed. Then a novel algorithm termed ‘local progressive abundance filter’ is applied to remove potential inter-genomic repeats, strain variability, and complex error patterns. The resulting minimizer-space contigs (mContigs) are added to the set of mReads for the next iteration. At the end of the multi-*k*′ process, reads are mapped to the final mContigs in order to output their polished sequences in base-space. In the following sections, we describe in more detail each of the major steps.

### 7.3 Multi-*k*′ minimizer-space de-Bruijn graph assembly

In classical de-Bruijn graph metagenome assembly, the choice of the *k*′-mer size is critical. Smaller *k*′-mers increase sensitivity as they recover overlaps between reads from rare species, and are less sensitive to sequencing errors. On the other hand, larger *k*′-mers yield higher-contiguity assemblies by resolving longer repeats. In order to get the best of both worlds, multi-*k* strategies have been introduced [23]. The assembler typically iterates over *k* values from values *k_min_* to *k_max_* by fixed increments. In each iteration, a de-Bruijn graph is constructed from the input reads and the contigs generated from the previous iteration.

In minimizer-space, there are three ways to increase the base-equivalent length of a *k*′-min-mer: decrease the density *d*, increase the minimizer length, or increase the value of *k*′. We rule out increasing the minimizer length *k* under the hypothesis that it would increase sensitivity to sequencing errors. Changing the density is in principle interesting as it only affects the distance between consecutive minimizers, however, it would require the recomputation of all minimizers within the mReads and mContigs for each iteration of the assembler, which would be computationally costly. Therefore, we decided to only increase the *k*′ parameter, the length of the *k*′-min-mer, as it does not require minimizers to be recomputed.

In metaMDBG we iterate over *k*′ from values 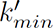 to 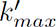 by increments of 1 (see section ‘Choice of parameters’ for the values and a discussion of (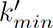, 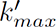)). The input reads are parsed only once to generate mReads, using fixed minimizer *k* and *d* parameters. Each iteration then extracts *k*′-min-mers from the mReads. Another advantage of this approach is that the base-space sequence of the contigs never needs to be constructed during the intermediate iterations: only the union of mContigs and mReads is used to construct the next graph.

### 7.4 Estimating *k*′-min-mers abundance and filtering errors

This part aims to refine the abundance of each *k*′-min-mer, i.e. the number of times a *k*′-min-mer is seen in the input reads. Generally, abundance information is used in de-Bruijn graph assemblers to detect and filter out erroneous *k*′-mers before graph construction, to reduce its complexity and memory consumption. Here the same philosophy is adapted and further elaborated for *k*′-min-mers. Refined abundances are estimated in two steps. 1) Prior to the first graph construction, *k*′-min-mer abundances are collected from raw *k*′-min-mer counts in mReads. 2) At each *k*′ iteration after graph construction, long mContigs, which are unlikely to be erroneous, are examined to refine the abundances of *k*′-min-mers and better detect erroneous *k*′-min-mers. Refined abundances are then propagated to the *k*′-min-mers of the next multi-*k*′ iteration.

#### Initial *k*′-min-mers counting and filtering

Even though the MDBG is a lightweight data structure, inserting all erroneous *k*′-min-mers would dramatically increase graph memory consumption and its complexity, making its traversal computationally challenging. Prior to constructing the graph for the first value of *k*′, we thus apply a abundance-based filter on *k*′-min-mers to remove the majority of erroneous ones. In metagenomics, detecting erroneous *k*′-min-mers is non-trivial, as low-frequency *k*′-min-mers might either correspond to real genomic sequences coming from rare species, or to errors. Our idea in this first step is to consider the *k*′-min-mers in the context of the read they have been extracted from: an estimate of a long read ‘abundance’ is determined, then its *k*′-min-mers having very low ‘local’ abundance are filtered out.

More precisely, we first perform *k*′-min-mer counting, similarly to classical *k*′-mer counting, i.e. the number of occurrences of each distinct *k*′-min-mer is determined. Then each read is processed sequentially. We define the read coverage *R_cov_* as the median of abundances of all its constituent *k*′-min-mers. We then determine a minimum abundance cutoff *R_min_* = *R_cov_ ∗ β* (where *β* = 0.1). A *k*′-min-mer is discarded if the following two criteria are satisfied: its abundance is equal to one, and is also lower than *R_min_*. This only removes *k*′-min-mers seen once, which represents the vast majority of erroneous *k*′-min-mers, but only within reads where *R_cov_*is greater than 1*/β*. Other potentially erroneous *k*′-min-mers will be detected during the contig generation process by the ‘local progressive abundance filter’ method described in the next subsection.

#### Refining *k*′-min-mer abundances

After mContigs have been generated (next section), *k*′-min-mer abundances are refined. We introduce two techniques: abundance smoothing and long contig *k*′-min-mer rescuing. The smoothing step is performed first. The abundance of a mContig *C_cov_* is computed as the median abundance of its constituent *k*′-min-mers. In the mContig, the abundance of each *k*′-min-mer is then set to the refined abundance *C_cov_*. Long mContigs (having *>* 2*k*′ k’-min-mers) are unlikely to contain any erroneous *k*′-minmers. If a *k*′-min-mer with abundance 1 is present in a long mContig, it is rescued by incrementing its refined abundance by 1 so that it will pass the pre-filtering performed in the next iteration.

#### Propagating refined abundance to the next *k*′ iteration and filtering

At the beginning of each sub-sequent multi-*k*′ iteration except the initial one (*k*′ > 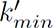), we estimate *k*′-min-mers based on the refined abundance of (*k*′-1)-min-mers determined in the previous iteration. A *k*′-min-mer contains two overlapping (*k*′-1)-min-mers for which the refined abundance is known. We define the refined abundance of a *k*′-min-mer as the minimum of its two (*k*′-1)-min-mers abundances. We use the minimum instead of the average because if one of the two (*k*′-1)-min-mers is erroneous, we do not wish its abundance to be raised by the other potentially correct one. This refined abundance propagation technique is to the best of our knowledge novel and has several advantages. Firstly, it improves *k*′-min-mer abundance estimation over using abundances determined from reads alone. Secondly, it prevents *k*′-min-mer abundances from collapsing to one (or even zero) as we increase *k*′: indeed long *k*′-min-mers tend to be underrepresented as they are more likely to contain a sequencing error or be longer than the mReads themselves. Finally, refined abundances allow us to assign an abundance estimate to *k*′-min-mers that only exist in mContigs and not in mReads.

After the *k*′-min-mer refined abundances have been determined, all *k*′-min-mers seen once are discarded. As we progress in the multi-*k*′ process, we notice that the refined abundances of erroneous *k*′-min-mers tends to become one, whereas correct *k*′-min-mers tend to get rescued and refined to abundances of two or more.

### 7.5 Local progressive abundance filtering

In this section, we introduce a key component of our contig generation process, which performs progressive abundance filtering to simplify parts of the assembly graph corresponding to abundant organisms (typically above 10-20× coverage). We first explain the rationale then give algorithmic details in the next paragraphs.

We generate contigs by examining the abundances of organisms in the assembly graph through the abundances of unitigs. Recall that a unitig is a maximal-length non-branching path in the assembly graph. Nearly all unitigs of abundant organisms cluster together into a single large connected component of the assembly graph. This is due to inter-genomic repeats and chimeric reads in HiFi samples. These two effects drastically increase the complexity of the graph and make assembly challenging. By performing graph simplifications using abundance information, we will sidestep both issues.

In principle, some abundant organisms could be separated *in silico* from the large component of the assembly graph using an abundance filter, e.g. by removing all nodes with abundance lower than half of the organism’s abundance. This is because most of the erroneous overlaps have low coverage: chimeric reads are rare, and most inter-genomic repeats are spanned by rare species, so removing the corresponding low-abundance graph nodes will remove those repeats. Filtering using a local abundance criteria has additional advantages: it can get rid of large stretches of sequencing errors, and can remove strain variability. However, designing such a filter is not straightforward.

In complex areas of an assembly graph unitigs tend to be fragmented and their abundances under-estimated, resulting in correct unitigs being filtered out whenever removal is based on their length, or more critically, their absolute abundance. Interestingly, the abundances of chimeric or rare species unitigs in complex areas also tend to be under-estimated [23]. Our remedy will be to filter out unitigs by iterating over abundance cutoffs, from low to high. At some point in the iterative process, fragmented but correct unitigs will be linked to longer ones, and thus successfully rescued.

An unpractical but simple algorithm that illustrates our contig generation process is as follows. Sort the MDBG unitigs *u*_1_, …, *u_n_* from the most abundant (*u*_1_) to the least abundant (*u_n_*). Iterate the following procedure from *i* = 1 *… n*. Consider the abundance *a_i_* of *u_i_* and fix a local abundance cutoff *U_i,cut_* = *a_i_ ∗ β* (with *β* values in the range 0.1 – 0.5, which in the real algorithm we will set it to 0.5). Create a copy *G*’ of the MDBG. For *t* = 1 to *t* = *U_i,cut_*, repeat removing all unitigs having abundance below *t* from *G*’ and re-compact *G*’. Finally at *t* = *U_i,cut_*, collect the unitig *u*’ in *G*’ that contains *u*. If *u*’ does not contain any *k*′-min-mer from a previously returned contig, return it as a contig.

Performing assembly with the above procedure for every unitig would be costly and redundant. Instead, in this work a progressive abundance filter is applied once to the whole graph from thresholds *t* = 1 to *t* = *t_max_*(see paragraph ‘Progressive abundance filtering’) instead of performing it per unitig. At each step we collect the set of unitigs from the graph. This results in multiple sets of unitigs (*S*_1_*, …, S_tmax_*), each corresponding to a single threshold *t*. A subsequent algorithm iterates over the *S_t_*’s and non-redundantly outputs all the unitigs that are above a well-chosen abundance threshold at each step (paragraph ‘Generating mContigs’).

#### Progressive abundance filtering

This process (Algorithm 1) iterates over abundance thresholds, simplifying and compacting the graph and then removing unitigs below the current threshold, saving the remaining unitigs.

Specifically the algorithm iterates from abundance threshold *t* = 1 to *t* = *t_max_* (line 3), where *t_max_* is the abundance of the most abundant unitig in the initial graph. The graph is simplified (line 4, see next section ‘Graph simplification’ for details). The graph is then compacted (line 5) and unitigs are collected into a set *S_t_* (line 6). Finally, unitigs with abundance *≤ t* are discarded (line 7) from the graph and we move to the next iteration of *t*.

##### Algorithm 1 Progressive abundance filtering.

**Input:** MDBG G

**Output:** *S*_1_*, …, S_t__max_* sets of unitigs along with their abundance information

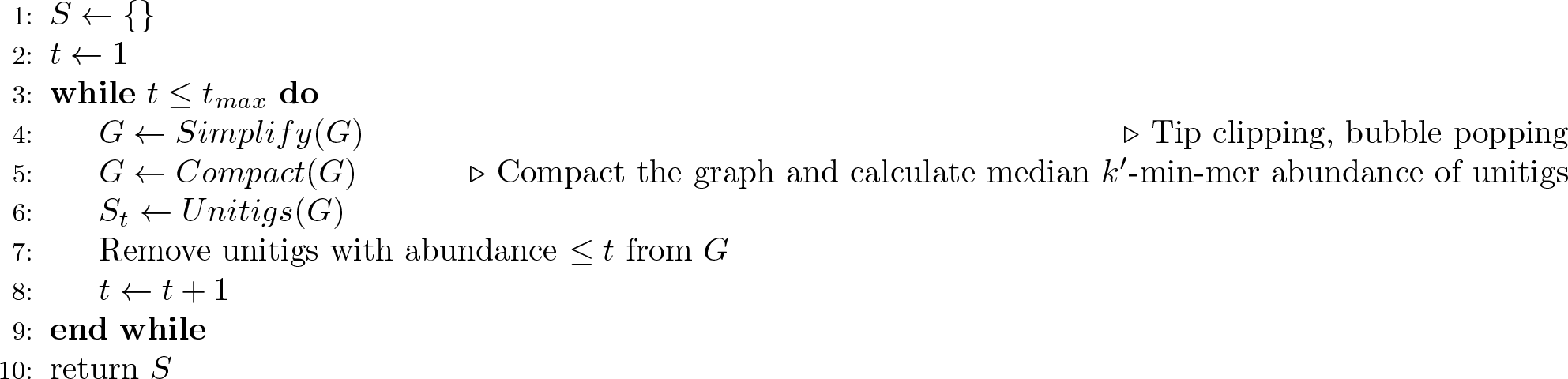

#### Graph simplification

The simplification step includes two processes: tip clipping and superbubble popping. Tips of 50 kbp or smaller are disconnected from the graph. We do not remove them here as they may either be erroneous or belong to a rare species. These tips are removed at the end of the assembly process if they have a high identity with another contig. Superbubbles of length 50 kbp or smaller are detected in *O*(*|Edges|*+*|Nodes|*) average time following the definition and algorithm of Onodera *et al* [24], and the path with maximum abundance is kept.

#### Generating mContig

This process iterates over all sets of unitigs *S_t_*’s starting from the one with highest abundance cutoff, *S_t__max_*. For each set, unitigs and their abundances are scanned in no particular order and an unitig *u* is returned if its abundance *a* is greater than some threshold. We call mContigs the set of returned unitigs (in line with typical genome assembly usage, where a contig is generally a unitig within the simplified assembly graph). The complete process is described in Algorithm 2.

Specifically at each iteration, a unitig *u* from *S_t_* along with its abundance *a* is added to the final set of mContigs if it does not share any *k*′-min-mer with any other unitig already in mContigs, and also if its abundance *a* is greater than *a ∗ t/β* (line 6). The *k*′-min-mers within *u* are recorded in a set of outputted nodes to prevent redundancy (lines 7 - 8).

Here the sets of unitigs *S_t_*’s are iterated from the large abundance threshold to the low abundance threshold, rather than the opposite. This is to make sure that we always output unitigs in their longest possible form. To see this, consider what would happen if we had started with the lowest threshold. There would be no way of knowing if a given unitig has been maximally merged with some other unitig(s) after our abundance filtering and graph simplifications steps. For example, at the abundance threshold of 3, all unitigs with an abundance of 6 would be output as they pass the local abundance threshold of 3*/*0.5 = 6. But among them there may also be fragmented unitigs belonging to a more abundant species (e.g. of abundance 10) that are ‘waiting’ to be merged with other unitigs after more drastic simplifications (at t=4 or 5 for instance). Iterating from the large threshold to the low threshold solves this.

### 7.6 Converting to base-space and assembly post-processing

At the end of the multi-*k*′ process the base-space representation of mContigs, i.e. the actual nucleotide sequences and not their minimizer-space representation, is constructed by gathering the base sequences corresponding to all mContigs *k*′-min-mers from the original reads. It is followed by two post-processing steps. A contig polishing step fixes sequencing errors in contigs (mostly homopolymers), and an optional duplication purging step removes similar contigs corresponding to close strains.

#### Constructing contig base sequences

This step converts mContigs, i.e. the minimizer-space representation of contigs, to actual nucleotide-space contigs. The idea is to choose a particular *k*′ value, collect *k*′-min-mers nucleotide sequences from the original reads, then reconstruct contig nucleotide sequences by aggregating the *k*′-min-mers nucleotide sequences. This is a generalization of the method presented in [14] to the multi-*k*′ setting, also made more accurate using read mapping. Indeed a *k*′-min-mer can be generated by multiple different nucleotide sequences. Hence collecting the ‘wrong’ nucleotide sequence could yield errors in contigs. Large values of *k*′ yield more specific *k*′-min-mers, minimizing such errors. However, some of these long *k*′-minmers may only exist in mContigs but not in mReads, thus their nucleotide sequences cannot be constructed with certainty. We use *k*′ = *k*′ to ensure that all contigs *k*′-min-mer are indeed present in the reads. To collect the ‘true’ nucleotide sequence of each contig *k*′-min-mers, mReads are first mapped to mContigs. *k*′-min-mers sequence are then collected from the reads that best match on contigs. The read mapping strategy in minimizer-space is described as follow.

##### Algorithm 2 Generating mContigs.

**Input:** *S*_1_*, …, S_t_*_max_ sets of unitigs

**Output:** mContigs

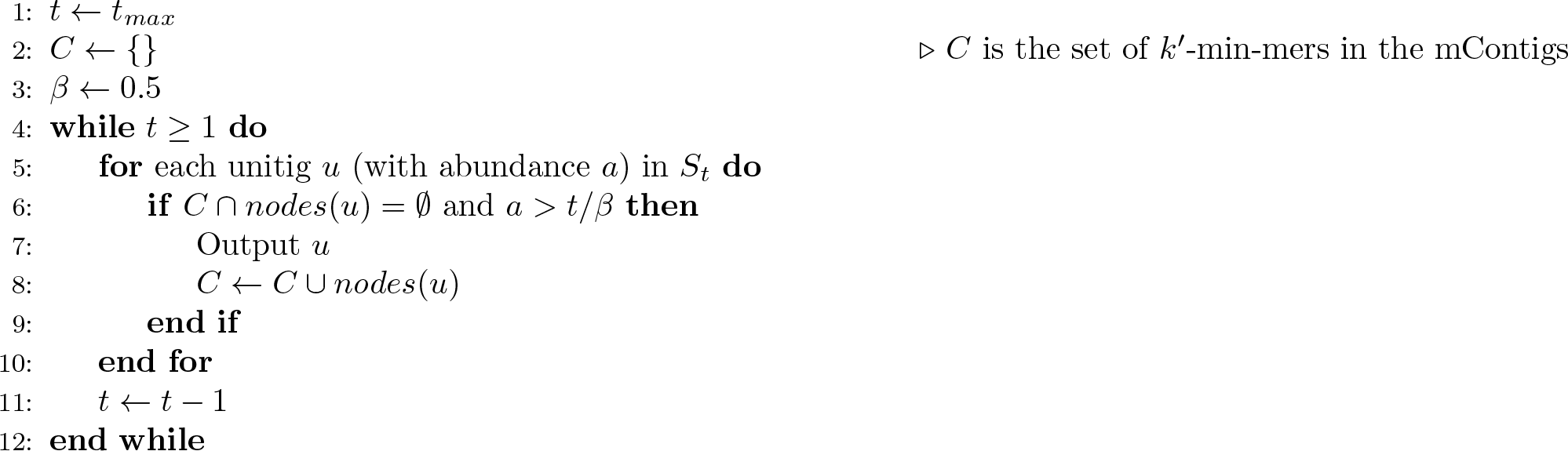

The mContigs are firstly indexed to create a set of *k*′-min-mer seeds: each mContig *k*′-min-mer is stored as a key in a hash table with the associated values being a list of contig *positions*, represented as pairs *{c_i_*, *c_p_}*, where *c_i_* is the contig identifier and *c_p_*is the *k*′-min-mer position in *c_i_*. Then mReads are scanned, and for each mRead *k*′-min-mer found in one or more mContigs, its mContig position(s) are retrieved as *seeds* for potential mappings. The seeds are extended maximally: we iterate over the mRead *k*′-min-mers (to the left and to the right of the seed) and extend mappings as long as subsequent *k*′-min-mers continue to be the same as those that follow in the mContig(s). The result is a set of intervals (made non-redundant) indicating maximal matches between the current mRead and one or several mContigs. Then another hash table with contig *k*′-min-mer positions *{c_i_*, *c_p_}* as keys (here *c_p_*is the position of the seed in the mContig, one position per mapping obtained), maintains the maximal matches as triplets *{r_i_, r_p_, m}* where *r_i_* is the read identifier, *r_p_* is the position of the seed *k*′-min-mer in *r_i_* and *m* is the length of the longest match.

The overall mapping algorithm is thus quadratic over the number of *k*′-min-mers in each mRead. However, in practice this number is close to 45, making the algorithm highly practical. We process mReads twice, in forward and reverse order, to handle reverse complements. The output of the algorithm is exactly one read *k*′-min-mer position for each contig *k*′-min-mer position.

The reads are then parsed in nucleotide space, and their *k*′-min-mers are extracted. If a *k*′-min-mer is one that was reported as a best match during the above mapping procedure, we collect the substring of the read corresponding to that *k*′-min-mer. To deal with overlaps between successive *k*′-min-mers in mContigs, we also record the position of the second and second-to-last minimizers within each *k*′-min-mer. We finally parse mContigs and concatenate the sequences associated to their *k*′-min-mers, making sure to discard overlaps.

#### Contig polishing

We perform an additional polishing step on the base-level representation of contigs to remove sequencing errors. We re-implemented a strategy akin to `racon` [25]: reads are first uniquely assigned to contigs using `minimap2`, contigs are then split into non-overlapping windows of 500 nucleotides and fragments of reads that map to each window are collected. Finally a consensus sequence for each window is created by partial order alignment using the SPOA library [25].

Our polishing differs from racon, in particular in the following two aspects. The first is how we select reads in case of multiple mappings. We noticed that longer alignments are not necessarily the best ones, but that alignment identity must also be considered. We score alignments using the metric *MS* = *alignLength ∗ alignIdentity* and only retain for each read the alignment that maximizes *MS*. The second is a reduction of memory usage: we limit the number of read fragments used to correct a window, with accurate long-reads, we noticed that using only 20 fragments is sufficient to produce a high-quality consensus; we also reduce the memory required to store the read fragments by partitioning the contigs and the reads that map onto them on the disk, processing one partition at a time. The memory required to store the read fragments of a contig is estimated by multiplying the contig length by the contig coverage (estimated from the initial read mapping). Contigs are processed sequentially and written into a partition file until the memory required to process the partition exceeds 6 GB. The current partition is then closed and a new one started. A structure in memory records the association of contigs to partitions. Similarly, reads are then processed and written to the partition of their best matching contig. This resulted in an approximately one-hundred fold reduction in memory usage compared to the original racon implementation for the Sheep rumen data set.

#### Strain duplication purging

Sequence duplications in contigs due to strain variablity are detected by an all-versus-all contig mapping using wfmash [26]. Contigs longer than 1 Mbp are left untouched and are used as templates to remove duplications present in shorter contigs. For those shorter contigs, we remove any part overlapping with a *≥* 1 Mbp contig when the overlap nucleotide alignment identity is over 99%.

### 7.7 Choice of parameters

Our method has four critical parameters: the minimizer size, the minimizer density, the starting and ending *k*′-min-mer size 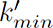 and 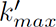.

The minimizer size and density were both set empirically to respectively 13 and 0.005 (*i.e.* rougly 0.5% of total *k*′-mers are used as minimizers). In our tests, using such short minimizers leads to superior results than using longer minimizers, possibly because they are less sensitive to sequencing errors.

The starting *k*′-min-mer size k’_*min*_ was fixed to 4. Using *k*′ values less than 4 creates assembly graphs that have high complexity, resulting in highly fragmented contigs. The ending *k*′-min-mer size 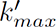 is a function of the sample median read length: 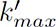 = *medianReadLength ∗ density ∗* 2.

With density 0.005 and 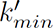 = 4, the assembler initially considers overlaps between reads of lengths 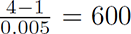 bases on average. It then iteratively increases the overlap length, in increments of 200 bases, until finally processing overlaps of twice the median length of the reads.

### 7.8 AD sample extraction and long-read DNA sequencing

Three biomass samples were taken at weeks 1, 20, and 40 from a year long sampling campaign, directly from an AD reactor digesting food waste by the facility operators and shipped in ice-cooled containers to the University of Warwick. Upon receipt, they were stored at 4*^◦^*C and then sampled into several 1-5mL aliquots within a few days and stored in 1.8 mL Cryovials at -80*^◦^*C. Samples were defrosted at 4*^◦^*C overnight prior to DNA extraction. DNA was extracted from a starting mass of 250 mg of anaerobic digester sludge using the MP Biomedical FastDNA SPIN Kit for Soil (cat no: 116560200) and a modified manufacturers protocol (see [27] for detailed protocol).

DNA size was assessed using a FemtoPulse (Agilent). The Pacific Biosciences protocol ‘Preparing 10 kb Library Using SMRTbell ®Express Template Prep Kit 2.0 for Metagenomics Shotgun Sequencing’ was used to create libraries from 1.5 micrograms of DNA. In most cases the DNA was already 10 kb or smaller. Sample AD2W40 was a bit larger so the DNA was sheared using a g-TUBE (Covaris) for one library and unsheared for a second library. Libraries were not pooled due to the large number of reads desired. Sequencing was performed using a Sequel II sequencer (Pacific Biosciences) using version 8M SMRT cells and version 2.0 sequencing reagents with 30-hour movies with 2 hr pre-extension time to generate CCS reads.

### 7.9 Assembling datasets, mapping reads and binning contigs

We ran all assemblers with 16 CPU threads. We used metaMDBG default parameters for all assemblies (minimizer size = 13 and density = 0.005). We ran hifiasm-meta with default parameters on real data, and with option –force-preovec on mock communities as suggested by the authors. We only used the hifiasm-meta primary assembly of polished contigs (p ctg.gfa), since adding alternate contigs reduced the overall MAG quality. We ran metaFlye with options –pacbio-hifi, –plasmids, –meta. We used the command ‘/usr/bin/time -v’ to obtain wall-clock runtime and peak memory usage. All tools used and complete command line instructions are available in Supplementary Table S1.

To determine the fraction of reads mapped to assemblies, we used ‘minimap2 -x asm20’ as suggested in the metaFlye study [6]. We filtered out a read when all its alignments were shorter than 80% of its length, and we assigned each remaining read to a unique contig through its longest alignment (breaking ties arbitrarily). To estimate contig coverage across samples prior to binning, we used the command ‘minimap2 -ak19 -w10 -I10G -g5k -r2k –lj-min-ratio 0.5 -A2 -B5 -O5,56 -E4,1 -z400,50’ as proposed in the hifiasm-meta article. We input the resulting BAM to the program `jgi_summa_rsize_bam_contig_depths` of MetaBAT2 to obtain contig coverage profiles across samples.

We performed contig binning using MetaBAT2 [28] with default parameters and a fixed seed (–seed 42) for reproducibility. Since MetaBAT2 may bin strains from the same species, creating a single apparently contaminated MAG, we separated all circular contigs of 1 Mb or longer prior to binning the remaining contigs, as suggested in the hifiasm-meta study [10].

### 7.10 Quality assessment of assemblies

We used CheckM (v1.1.3) to assess the quality of all MAGs and circular contigs longer than 1 Mbp. We used viralVerify [17] (v1.1) to identify plasmids and virus in each assembly (Supplementary Table S2). We only considered contigs shorter than 500 kbp with prediction score higher than 5. Annotations labeled as ‘Plasmid’ or ‘Uncertain - plasmid or chromosomal’ were considered as plasmids, and similarly annotations labeled as ‘Virus’ or ‘Uncertain - viral or bacterial’ were considered as virus. We used Barrnap (https://github.com/tseemann/barrnap), and Infernal [29] to predict respectively rRNA and tRNA genes from circular contigs. We filtered out annotations with E-value over 0.01. A total of 437 (96%) near-complete circular contigs found by metaMDBG have one copy of the 5S, 16S and 23S genes and at least 18 tRNA genes, against 96.6% for hifiasm-meta, and 98.5% for metaFlye (Supplementary File S2).

### 7.11 Assessment of completeness and fragmentation of assemblies with reference sequences

We used the following process to assess the completeness and fragmentation of assemblies when reference genomes are available (mock reference genomes or near-complete circular contigs). We used wfmash to align contigs against the reference sequences. Alignments with less than 99% identity were filtered out. Alignments were ordered by their matching score *MS* = *alignLength ∗ alignIdentity* (best score first). We considered alignment identity to improve contig assignment to similar strains. Alignments were then processed sequentially and contigs were uniquely assigned to references. During this process, we check whether a reference is complete or not, meaning that at least 99% of its positions are covered by contigs. We prevent other contigs from being assigned to a complete reference. Moreover, we prevent a contig to be assigned to a reference if more than 30% of its matching positions are already covered by another contig. In this case, we first try to assign this contig to another reference. References with less than 70% completeness were considered missed by the assembler.

### 7.12 Taxonomic classification of MAGs recovered from AD samples

The phylogenetic tree of Figure 3 was built using fasttree [30] from the output alignment of gtdbtk version 2.1.0 [31] on near-complete quality MAGs of all three assemblers for the anaerobic digester dataset. Concurrent diversity coverage between the different assemblers was explored at different taxonomic levels from genus to domain. To do so, it is necessary to first address MAGs for which no annotation is available at a given taxonomic rank. A pair of unannotated MAGs may or may not share the same taxa. A first pass based on tree topology allows us to select neighbouring MAGs as candidates for sharing the same unknown taxa. As a second step we compute the Relative Evolutionary Distance using the R library Castor version 1.7.3 [32]. Following guidelines from gtdb, we use their median RED values for each taxa in order to decide on grouping together unknown MAGs. We then find the best ancestor for each unknown MAG in terms of their RED being nearest to the corresponding taxa median RED. If they share the same best ancestor, we group them together otherwise we split them into distinct unknown taxa. Tree manipulation and representation is carried out using the library ggtree version 2.4.1 [33], treeio version 1.14.3 [34] and ggtreeExtra version 1.0.2 [35].

## 8 Assembler implementation details

During transformation to minimizer-space, reads are homopolymer-compressed [36]. We handle reverse recomplements in a similar but different manner to classical de Bruijn graph assembly. We considered canonical *k*′-min-mers in the following manner: a *k*′-min-mer is compared to its reverse (and not its reverse complement). The first minimizer of each is compared: the *k*′-min-mer with the smallest minimizer is selected as the canonical representative. In case of equality, the second minimizer of each is compared, and so on. Note that minimizers are also considered in their canonical representations, which is in this case identical to the classical technique: a minimizer is in canonical form if its forward sequence is lexicographically equal or smaller than its reverse-complement sequence.

## 9 Data availability

All datasets used is this study were downloaded from NCBI Sequence Read Archive (SRA) with accession numbers given in Table 1. Zymo mock reference genomes are available at https://s3.amazonaws.com/zymo-files/BioPool/D6331.refseq.zip. ATCC mock reference genomes are available at https://www.atcc.org/products/msa-1003.

## 10 Code availability

MetaMDBG is available at https://github.com/GaetanBenoitDev/metaMDBG. We have also made the analysis scripts used in this study to compare assemblers available at https://github.com/GaetanBenoitDev/MetaMDBG_Manuscript.

